# Deep learning-based risk stratification for HER2-negative breast cancer patients

**DOI:** 10.1101/2021.05.26.445720

**Authors:** Mustafa I. Jaber, Liudmila Beziaeva, Christopher W. Szeto, Stephen C. Benz

## Abstract

In this paper, we present our analysis of the tumor microenvironment in digital pathology images to stratify risk in HER2-negative breast cancer patients using clinicopathological, spatial image, and cell-based features in a Cox’s proportional hazard model. We start the analysis by processing a set of 304 training pathology images using our in-house pan-cancer trained tumor, stroma, and lymphocyte region identification convolutional neural networks. The next step is computation of spatial regions of interest, namely: lymphocytes within (and adjacent to) tumor, lymphocytes within (and adjacent to) stroma, and stroma within (and adjacent to) tumor areas. Various cell-level information in these regions are then summarized, augmented to clinicopathological data, and linked to patient’s survival to train a Cox’s proportional hazards model. The proposed model outperformed a baseline model based on clinicopathological features only in analysis of an untouched test set of 202 whole slide images with p 8.49E-08 (HR = 0.4273).

## INTRODUCTION

To establish a framework for stratifying risk in HER2-negative breast cancer patients, we analyzed tumor microenvironment (TME) [1] components in breast cancer histopathology images by use of a set of three in-house trained deep learning - convolutional neural networks (ConvNets) [2] for detecting tumor, stroma, and lymphocyte regions. We then use a spatial image profile that abstracts such TME information at image level, in addition to clinicopathological features, in a Cox’s proportional hazards model [3] to stratify risk up to 5 years after diagnosis. As reported in the literature, deep learning technology has been applied to histopathology images of one [4] or several [5, 6] cancer types to predict outcomes. In Bychkov *et al.* [4], two types of deep learning architectures - convolutional and recurrent - were used to predict colorectal cancer outcome based on digitized images of hematoxylin and eosin (H&E) stained diagnostic tissues only. In Zhu *et al.* [5], they suggested patients of the same tumor stage based on their clinical trial reports could be predicted to have different outcomes and linked tumor visual appearance to individual patient survival. They validated this hypothesis successfully in brain cancer and lung cancer cohorts. An unsupervised K-Means clustering algorithm was used in that report as a tool to select a reasonable and sufficient number of sample patches representing different TME components rather than aggregating results of all image patches in the final prediction result. A study that focused on analysis of brain tumors and predicting outcomes is found in Mobadersany et al. [7]; they used both genomic biomarkers and microscopic images of tissue biopsies in a deep learning computational approach to predict the overall survival of patients. The method used adaptive feedback to simultaneously learn visual patterns and molecular biomarkers associated with patient outcomes. A more comprehensive study of a variety of cancer types is presented in Wulczyn *et al.* [6], wherein they proposed a deep learning-based system to predict disease specific survival across ten cancer types and compared them to baseline models based on features such as stage, age, and sex.

In our Cox’s model, cell-level information is processed along with a proposed spatial image profile that captures TME image-level information for patient’s risk prediction. Here, basic cell-level information such as count, area, perimeter, circularity, and eccentricity, among others, is captured by off-the-shelf software [8] and augmented to the spatial image profile to further emphasize some TME details that affect risk stratification. It has been reported that several cellular morphological features are correlated to gene clusters. For example, in Wang *et al.* [9] they demonstrated that 4 cellular features were able to separate a cohort of triple negative breast cancer patients to sets with significantly different survival outcomes.

Another study that focused on early-stage estrogen receptor-positive breast cancer patients and used nuclear shape and orientation features in H&E images to predict overall survival is found in Lu *et al.* [10]. They developed an image-based system to identify breast cancer patients that have a high likelihood of benefitting from adjuvant chemotherapy.

As described in Malherbe *et al.* [1], the breast cancer microenvironment can be subdivided into three main subsets: local, regional, and distant. This is applied in our automated machine-vision system for characterizing TME at two different resolutions. The first is at the pixel-based level, which is modeled by the cell-based features, and the second is at the patch-based level, which is of 100 x 100 square micron (μm^2^). This patch size is used in our in-house tumor, stroma, and lymphocyte detection systems. Given that many cells would fit in a patch of this dimension (the average size of an epithelial cell is 25 μm), patch-level labelling of tumor (vs. nontumor, for example) guarantees that the majority of the cells within the patch are indeed tumor cells. In our system, a patch can have more than one label. For instance, a patch with dual labels, such as tumor and lymphocytes, could have ~50 % tumor cells and 50 % lymphocytes. A patch with this characteristic would be assigned to the tumor-infiltrating lymphocytes (iTILs) feature in the spatial image profile, as will be detailed later in the Methods and Results section. It is also possible for an image patch to be detected as tumor only and be located next to a patch that is detected as lymphocytes only. These adjacent patches would be assigned to the tumor-adjacent lymphocytes (iTALs) feature. In addition, stromal tumor-infiltrating lymphocytes (sTILs) image feature, described in Hudecek et al. [11], is also used in our system as a descriptor of tumor, stroma, and lymphocytes co-localizations.

## DATASET

TCGA breast cancer cohort was used in this study and specifically included patients with at least one diagnostic whole slide image (WSI) and available hormone receptor (HR) status (positive or negative). They were sorted into a training set (n = 304) and an untouched testing set (n = 202). No HER2-positive patients were included in this study. Note we used a single WSI per patient in our analysis.

## METHODS AND RESULTS

### Clinicopathology-based system

Several sets of features were used in a Cox’s proportional hazard model to predict risk of survival for new patients. The first set, which contains clinical features, is used to draw the baseline model. This set includes: HR status, ethnicity, race, primary diagnosis, prior malignancy, prior treatment, treatment type, tissue of origin, and, AJCC pathologic primary tumor (T), regional lymph nodes (N), and for distant metastasis (M). Note that AJCC pathologic stage was not included in the baseline model as a feature as it correlates with the AJCC pathologic T, N, and M.

The forest plot of clinicopathology-based groups resulting from the baseline Cox’s proportional hazard model is shown in Fig. 1a. The most predictive clinical features in this baseline model are prior treatment and ethnicity. The baseline Cox’s model was trained on training data (n = 304) as shown in Kaplan-Meier plot in Fig. 1b with p 1.05E-08 (HR = 0.4681) and tested on testing data (n = 202) with p 3.92E-04 (HR = 0.5586). Note that the average risk value (estimated by the baseline Cox’s model) over all patients in the training set is used as a threshold to label test patients as “low risk” vs “high risk”.

**Fig. 1.**
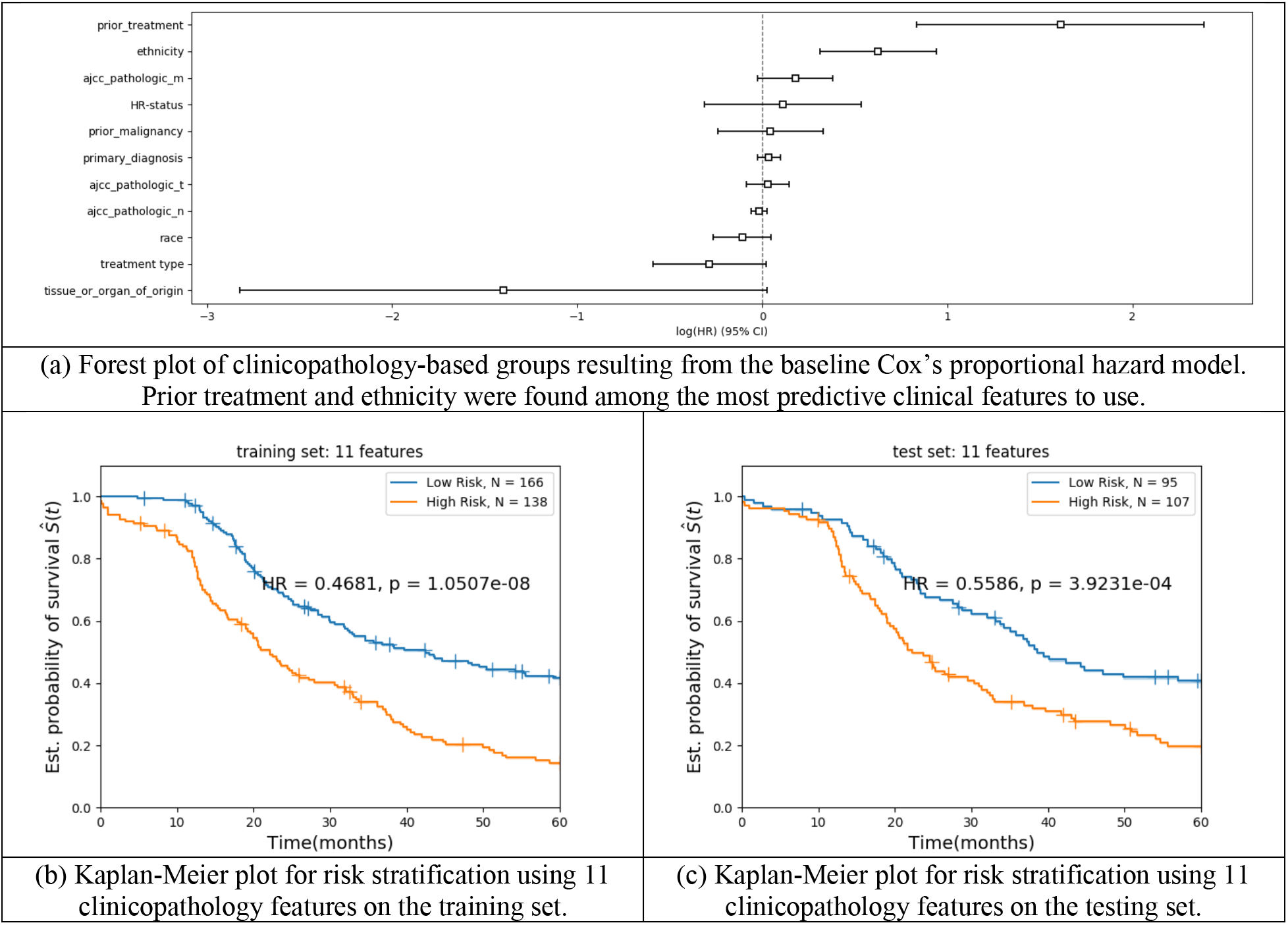
Cox’s proportional hazard model trained and tested using 11 clinicopathology-based features to form the baseline system for overall risk stratification.

Because HR status was a factor in selecting breast cancer patients for our training and testing cohorts, two Cox’s proportional hazard models were built (trained and tested) for HR positive and for HR negative sets, separately. Fig. 2 shows Kaplan-Meier plots based on these two Cox’s models for HR positive training (n = 255) and testing (n = 169) sets (in Fig. 2a and Fig. 2c) and HR negative training (n = 49) and testing (n = 33) sets (in Fig. 2b and Fig. 2d). The testing p values in Fig. 2c (p 1.61E-03) and in Fig. 2d (p 7.53E-02) are greater than the one in the baseline system in Fig.1c (p 3.92E-04) which indicate inferior performance. In the following discussion, we will use Cox’s model in Fig. 1 as the baseline model.

**Fig. 2.**
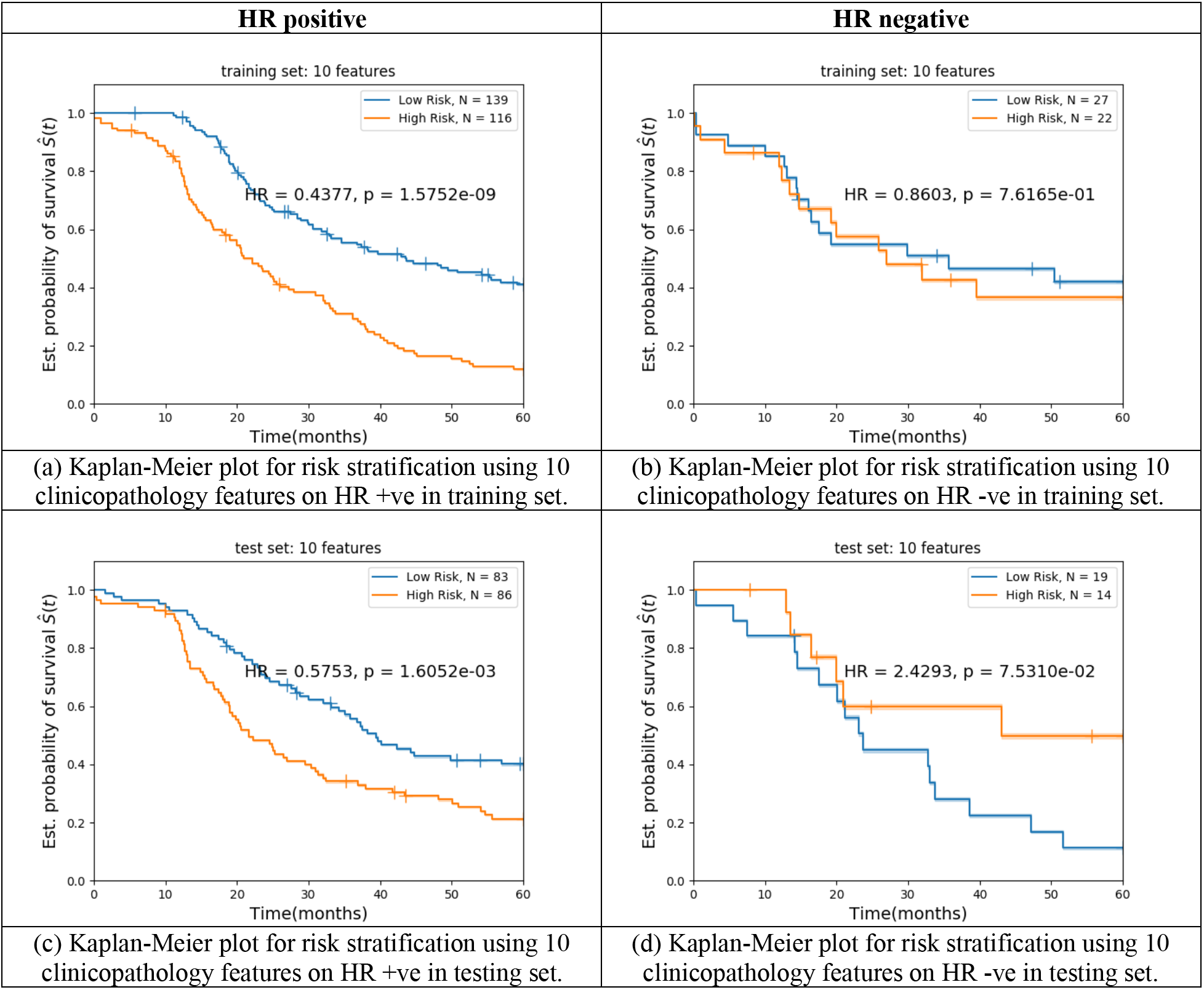
Cox’s proportional hazard model per hormone receptor status trained and tested using 10 clinicopathology-based features.

### Image-based system

We further add image-based spatial features to the baseline system. Features related to location, size, and co-occurrence of tumor, stroma, and lymphocytes regions are estimated using three different convolutional neural networks. The process starts by cutting input WSIs into square image patches which are then processed by three deep-learning image ConvNets to identify tumor, stroma, and lymphocytes regions. Next, the detected labels of neighboring image patches are used to build a spatial WSI profile which includes nine image features. Finally, these image features are combined with clinicopathology features to improve overall risk stratification.

In the following sub-section, we introduce pan-cancer tumor, stroma, and lymphocytes region detection modules and in the subsequent sub-section, we discuss the spatial image profile. Finally, we describe our overall risk stratification system for HER2-negative breast cancer patients based on clinicopathology and image features.

#### Pan-cancer tumor, stroma, and lymphocytes region detection modules

The human-in-the-loop annotation tool, shown in Fig. 3 and detailed in the patent “Few-shot learning based image recognition of whole slide images at tissue level” [12] was used to develop patch-level gold standard masks for tumor, stroma and lymphocytes regions. Sample WSIs from several TCGA cohorts were used as training sets.

**Fig. 3.**
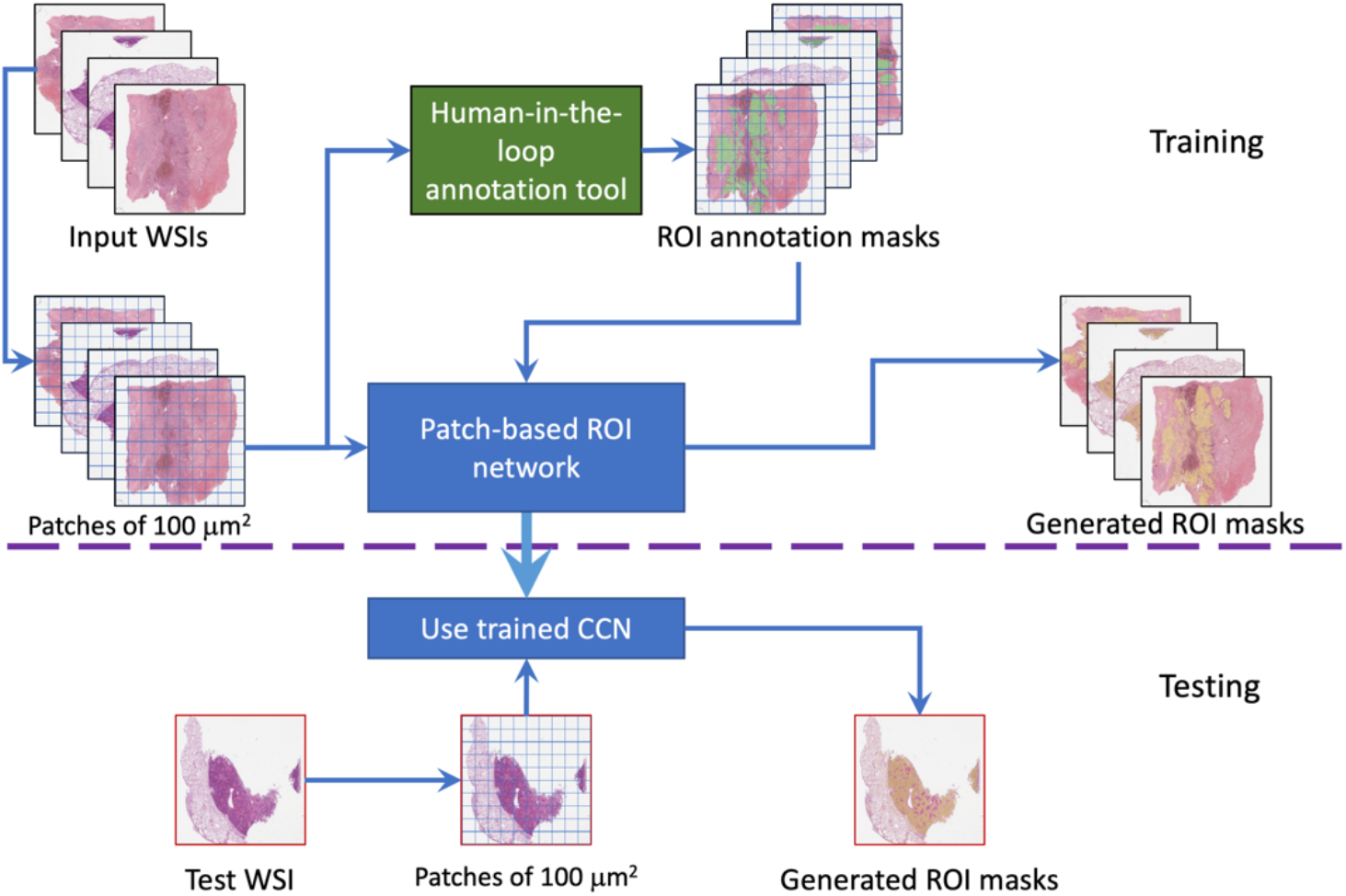
Block diagram of ROI identification. ROI could be tumor areas, stroma regions, or lymphocytes regions in a WSI. Human-in-the-loop annotation tool is from Song and Jaber patent [12].

Figure 3 shows two branches of WSI processing. First, TCGA WSIs used for training a supervised deep learning network are processed via the human-in-the-loop annotation tool to build corresponding patch-based level gold standard masks. The mask information and image patches are used to train a patch-based ROI identification ConvNet which is then used to autogenerate ROI masks. The generated masks are compared to the given gold standards for performance evaluation on training data. TCGA and internal WSIs without pathology annotations are then processed via the trained ConvNet for ROI identification. Here we introduce Inception-v3 architecture used for identification of tumor epithelium then we present the ResNet34 architecture used for lymphocytes region detection, and finally the ResNet34 architecture used for stroma detection.

##### i- Pan-cancer tumor identification

Detection of epithelial tumor regions in standard H&E stained histology images of different cancers is challenging due to the high variability and irregularity in size, shape, and chromatin texture of the tumor nuclei [13]. To handle such complexities, we trained a large deep learning architecture (Inception-v3).

The human-in-the-loop annotation tool presented in the Song and Jaber patent [12] was tuned to help pathologists annotate patch-level tumor masks for 433 WSIs from ten different TCGA cohorts. This set was further used to train an inception v3 tumor detection network as shown in Fig. 3.

##### ii- Pan-cancer lymphocytes region identification

A ResNet34 architecture was trained to detect lymphocytes regions in 80 TCGA diagnostic H&E-stained WSIs from different cancers. In this case, the human-in-the-loop annotation tool was tuned to help pathologists annotate patch-level lymphocytes.

##### iii- Pan-cancer stroma identification

A ResNet34 architecture was trained to detect stroma regions in diagnostic H&E-stained WSIs of nine different cancers. In our analysis, we consider anything other than tumor, normal, and lymphocytes regions to be stroma.

#### Spatial image profile

The spatial image profile in this study is consisted of nine features, namely:

1. **% tumor**: percent of tissue patches that are identified as tumor,
2. **% stroma**: percent of tissue patches that are identified as stroma,
3. **% lymphocytes**: percent of tissue patches that are identified as lymphocytes,
4. **% iTILs**: percent of lymphocytes patches within 100 μm from tumor (tumor-infiltrating lymphocytes),
5. **% iTALs**: percent of lymphocytes patches within 200 μm from tumor (tumor-adjacent lymphocytes),
6. **% sTILs**: percent of lymphocytes patches within 100 μm from stroma (stroma-infiltrating lymphocytes),
7. **% sTALs**: percent of lymphocytes patches within 200 μm from stroma (stroma-adjacent lymphocytes),
8. **% stroma in tumor**: percent of stroma patches within 100 μm from tumor patches, and
9. **% stroma near tumor**: percent of stroma patches within 200 μm from tumor patches.

Notice all features are in the range of 0 to 100.

Figure 4 shows a visual example of the generated masks and their spatial relations in a TCGA diagnostic WSI. The top row starts (from the left) with a WSI (Fig. 4a), followed by its generated tumor, stroma, and lymphocytes masks (Fig. 4b-d). The top row ends with the percentage of lymphocytes patches that co-locate with tumor (Fig. 4e) while the bottom row starts with the percent of lymphocytes adjacent to tumor area (Fig. 4f). Similar spatial relations are introduced for lymphocytes and stroma, that is, lymphocytes within stroma regions (Fig. 4g) and lymphocytes adjacent to stroma regions (Fig. 4h). Lastly, percentage of stroma in tumor regions (Fig. 4i), and percentage of stroma adjacent to tumor regions (Fig. 4j) are shown in the bottom row of Fig. 4.

**Fig. 4.**
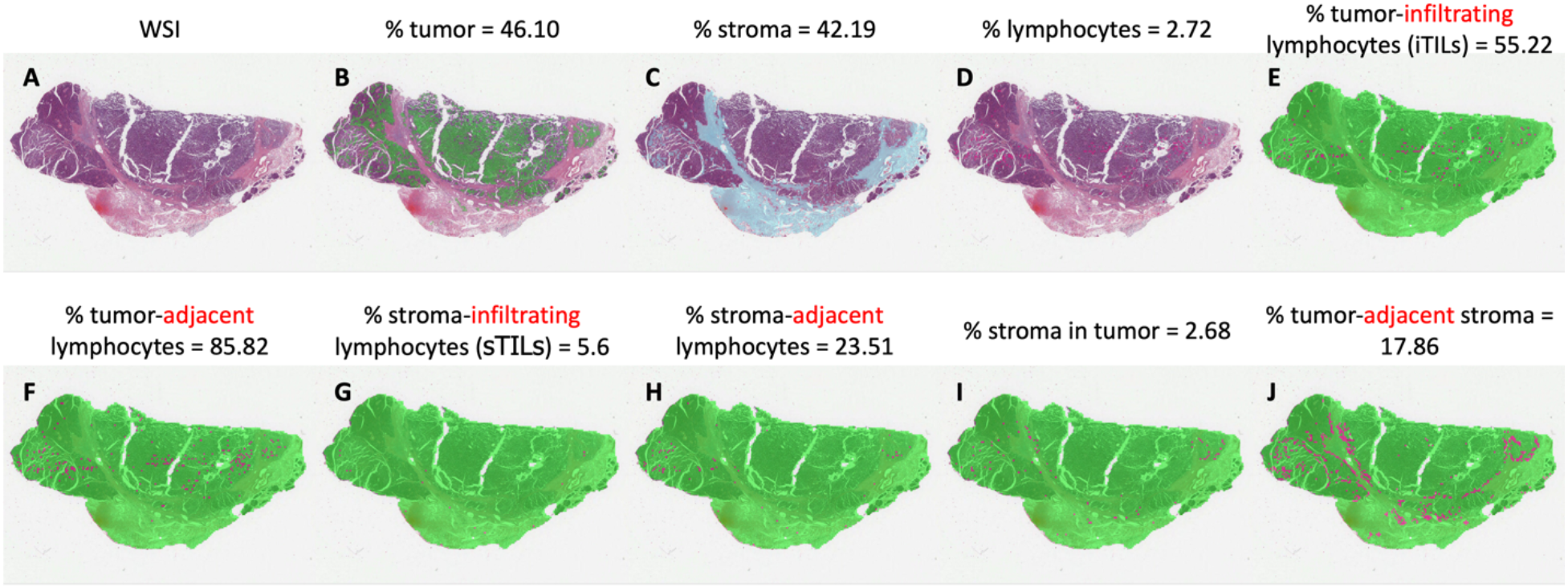
Spatial image profile. An image example of the nine spatial image features (based on auto-generated tumor, stroma, and lymphocytes masks) used in this study is presented. Green color in B represents tumor, sky blue color in C represents stroma, red color in D represents lymphocyte, green color in E-J represents tissue area, and red color in E-J represents mask area as given in the corresponding figure title.

Fig. 5 shows five visual examples of the developed auto-generated tumor, stroma, and lymphocytes masks. Other features of spatial image profiles are given as values. Examples of high and low tumor content are given (TCGA-AR-A24K-01Z-00-DX1 vs TCGA-E2-A15I-01Z-00-DX1). Similarly, examples of high and low lymphocytes content are also given (TCGA-BH-A0B9-01Z-00-DX1 vs TCGA-E2-A15I-01Z-00-DX1).

**Fig. 5.**
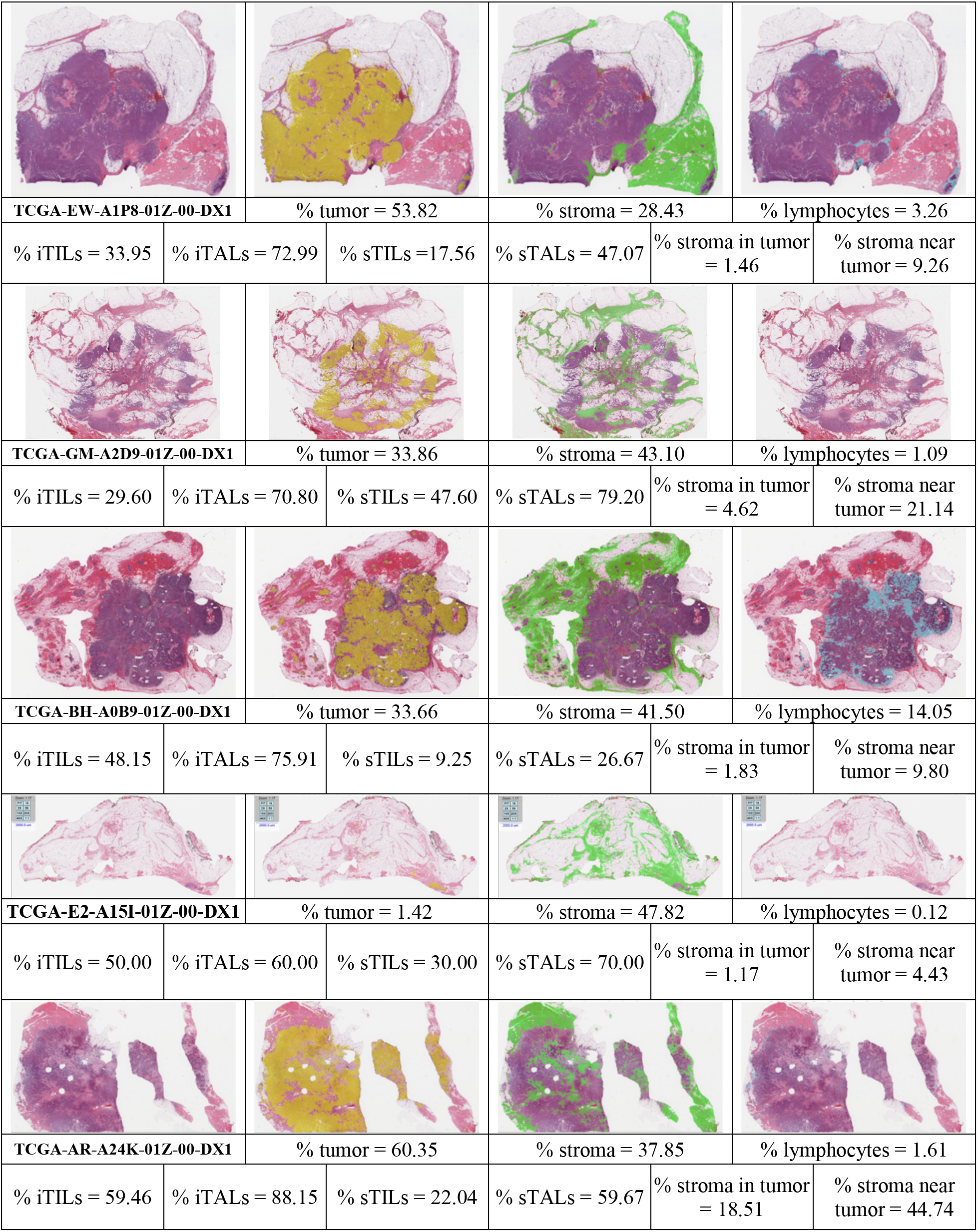
Five visual examples of auto-generated tumor, stroma, and lymphocytes masks. Other features of spatial image profiles are given as values. Golden yellow color represents tumor areas, green color represents stroma, and sky blue color represents lymphocytes regions.

#### Clinicopathology- and image-based system for risk stratification

To assess the accuracy of the model, a new Cox’s proportional hazard model was trained using 20 clinicopathology- and image-based features for overall risk stratification in HER2-negative breast cancer training set and tested on the untouched tested set is shown in Fig. 6. The performance was increased from p = 3.92E-04 (HR = 0.5586) on the baseline model to p = 1.31E-07 (HR = 0.4294) on this test set.

**Fig. 6.**
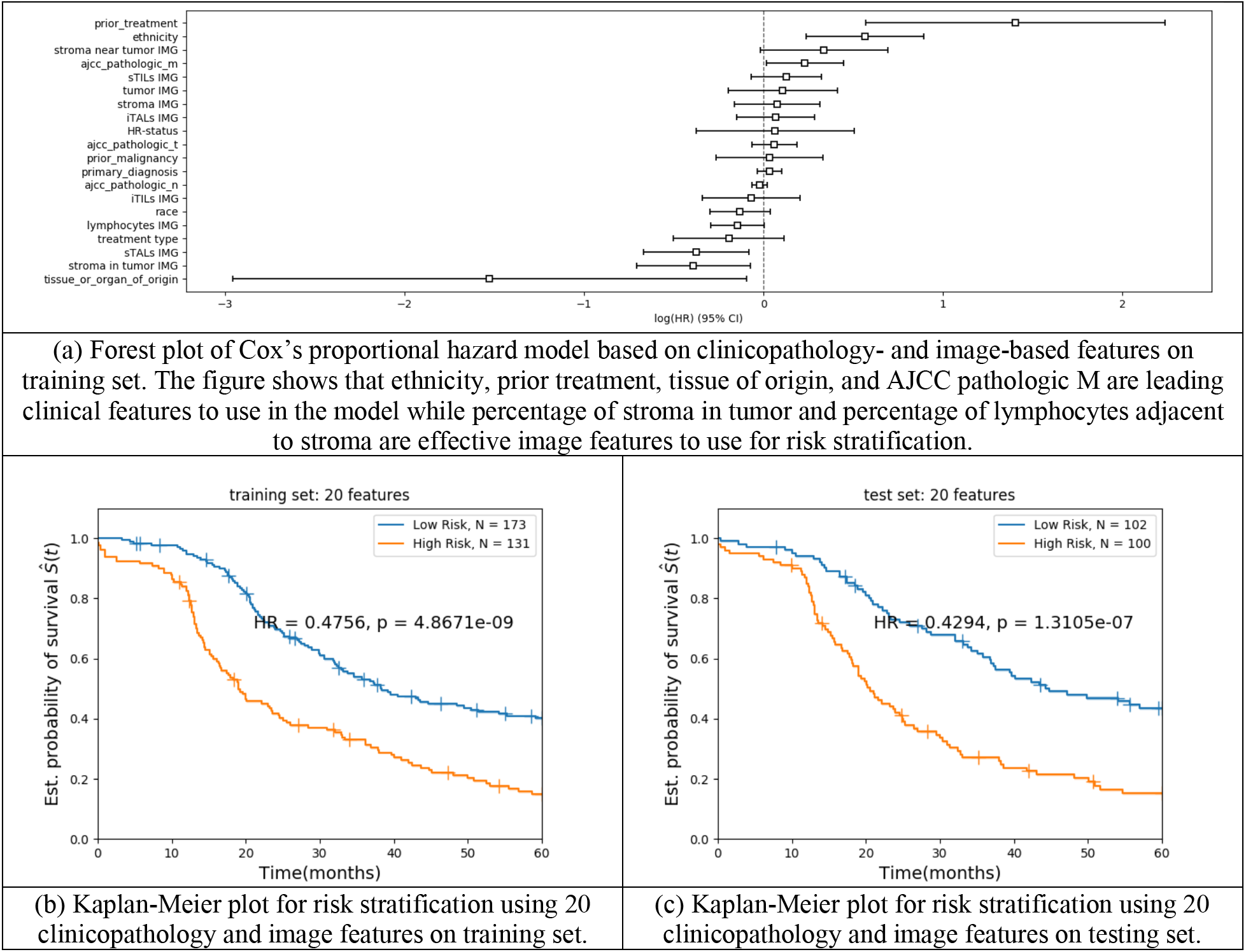
Cox’s proportional hazard model trained and tested using 20 clinicopathology- and image-based features for overall risk stratification in HER2-negative breast cancer patients.

Fig. 6a shows that when training and testing a Cox’s proportional hazard model based on clinicopathological and image features for overall risk stratification in HER2-negative breast cancer patients, the most important clinical features to use are ethnicity, prior treatment, tissue of origin, and AJCC pathologic M. In addition, percentage of stroma in the tumor and percentage of lymphocytes adjacent to stroma are also effective image features to use.

### Cell-based system

To study cell-level contributions to TME, we used off-the-shelf software in [8] to extract seven color- (and stain-) independent features such as nucleus count, nucleus distance, nucleus area, nucleus perimeter, nucleus max caliper, nucleus circularity, and nucleus eccentricity. In this system, these features were extracted per cell, then averaged per spatial image profile from previous section (when applicable). That is, all seven cell-based features were averaged for tumor regions, stroma regions, lymphocytes regions, iTILs regions, and so on. This process generates a total of 63 cell-based features per WSI.

#### Clinicopathology-, image-, and cell-based system for risk stratification

In order to combine features of different scales (different value ranges) into one model, quantile transformation with normal distribution was learned (on training data) and applied (on testing data) per feature. This preprocessing normalization step mapped all features to the same range. This transformation tends to spread out the most frequent values and reduces the impact of (marginal) outliers.

The Cox’s proportional hazard model trained using the updated 83 clinicopathology-, image-, and cell-based features for overall risk stratification is shown in Fig. 7 with HR = 0.4273 and p = 8.49E-08 on the testing set. Its performance is better than the baseline model (HR = 0.5586, p = 3.92E-04) and the clinicopathology and image model (HR = 0.4294, p = 1.31E-07) on the same test set. Fig. 7a shows a forest plot of the trained Cox’s proportional hazard model based on clinicopathology-, image-, and cell-based features of the training set. The figure shows that prior treatment and AJCC pathologic M are the most useful clinical features to apply in the model. It also shows that the percentage of lymphocytes adjacent to stroma is an effective image feature to use for risk stratification in the model. The data in the figure support the use of the following cell-based features in the model: nucleus max caliper and nucleus eccentricity in iTILs regions, nucleus area and nucleus max caliper in stroma-in-tumor regions, nucleus count in stroma-near-tumor regions, and nucleus max caliper and nucleus eccentricity in lymphocytes regions.

**Fig. 7.**
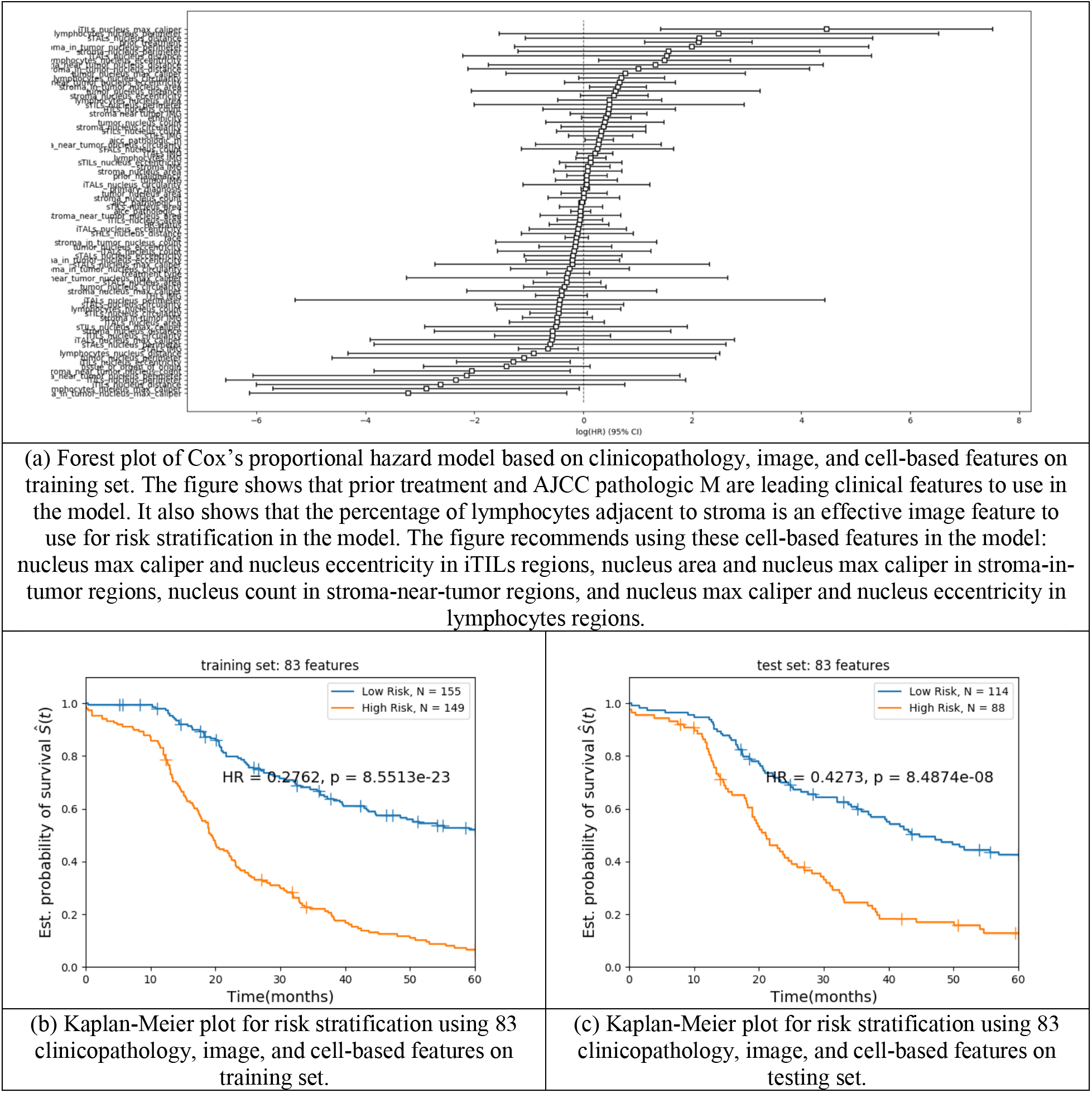
Cox’s proportional hazard model trained and tested using 83 clinicopathology, image, and cell-based features for overall risk stratification in HER2-negative breast cancer patients.

## DISCUSSION

### Hormone receptor status

Hormone receptor status was found to be a factor “not contributing” in all Cox’s proportional hazard models (Fig. 1a, Fig. 6a, and Fig. 7a) introduced in this study. To analyzed that further, we combined training and testing data and applied the risk prediction model in Fig. 7a to generate Fig. 8a. The predicted low risk group (n = 269) was further separated into two groups based on their hormone receptor status in Fig. 8b with hazard ratio of 1.1551 and p = 4.42E-01. Similarly, the predicted high risk group in Fig. 8a (N = 237) was separated into two groups based on their hormone receptor status in Fig. 8c with hazard ratio of 1.0970 and p = 5.85E-01. The resulting hazard ratio ~ 1 confirms that the two Kaplan-Meier curves in Fig. 8b (and the two in Fig. 8c) are statistically not separable. This indicates the developed risk stratification system is independent of hormone receptor status.

**Fig. 8.**
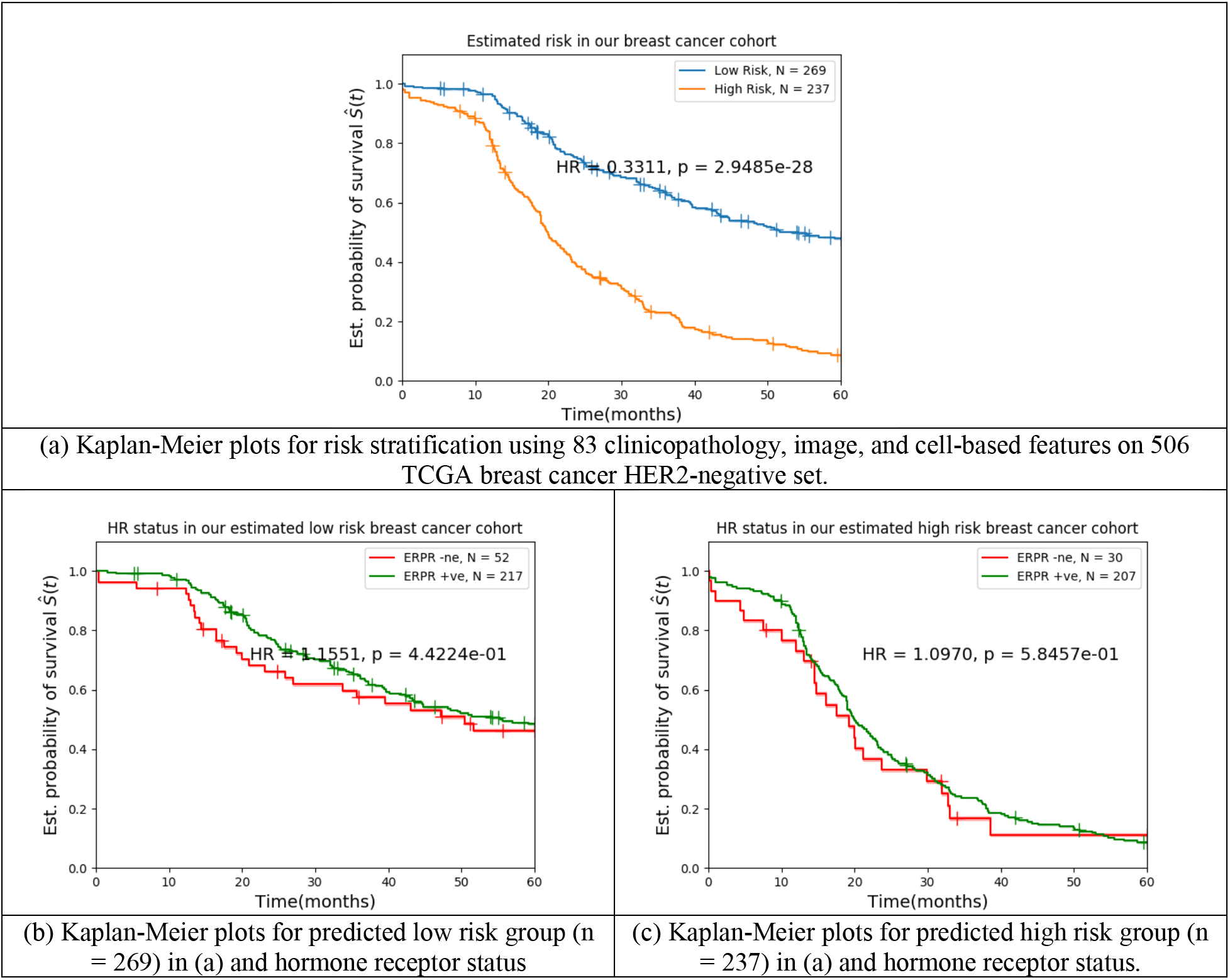
Hormone receptor status in HER2-negative breast cancer patients.

### Case study

Predicted risks based on five digital images shown in Fig. 5 from the test set are given in Table 1. The predictions from the Cox’s proportional hazard model for overall risk stratification in HER2-negative breast cancer patients presented are based on 1) clinicopathological features only (baseline model in Fig. 1a), 2) clinicopathological and image features (Cox’s model in Fig. 6a), and 3) clinicopathological, image, and cell-based features (Cox’s model in Fig. 7a). Time after diagnosis in months, status, and hormone receptors status are also shown. The most predictive features of these three Cox’s models for the WSIs in Table 1 are given in the supplementary section in Table S1, Table S2 and Table S3.

**Table 1.**
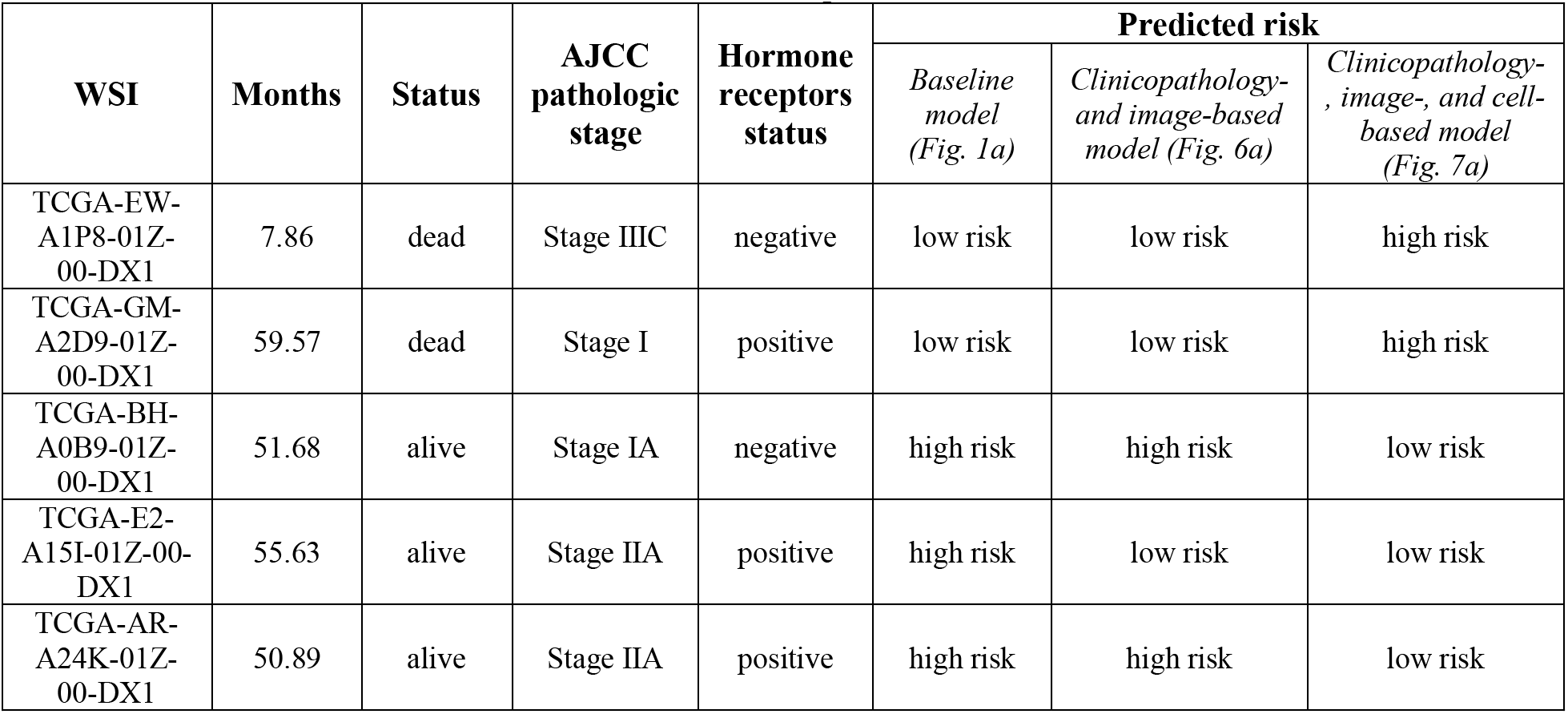
Predicted risks in various Cox’s proportional hazard models for overall risk stratification in HER2-negative breast cancer patients

For the given examples, the Cox’s model in Fig. 7a almost always gave more accurate risk predictions than the baseline model. For example, the baseline system predicted TCGA-EW-A1P8-01Z-00-DX1 who lived only 7.86 months after diagnosis as low risk which is incorrect; our Cox’s model in Fig. 7a more accurately predicted that as high risk case.

The baseline model predicted patient TCGA-GM-A2D9-01Z-00-DX1 as low risk, particularly given she lived for almost 60 months after diagnosis with breast cancer with AJCC pathologic Stage I (Note our study had a limit of 60 months survival after diagnosis as shown in all Kaplan Meier plots in this manuscript). In contrast, the Cox’s model in Fig. 7a identified sufficient details in the image- and cell- level features to predict the risk for this patient was high.

Conversely, the baseline system predicted the rest of the cases shown in Table 1 to be high risk, but all lived for more than 50 months after diagnosis, pointing to the low accuracy of that system. Our Cox’s model in Fig. 7a predicted low risk for all, which was found to be more accurate.

The clinicopathology- and image-based model that was introduced in Fig. 6a gave mixed prediction results.

## CONCLUSION

In our proposed framework for stratification of risk in HER2-negative breast cancer, automated machine-vision features that characterize the tumor microenvironment at image- and cell-levels are considered along with clinicopathological features in a Cox’s proportional hazard model. We demonstrate the developed image-level TME descriptors and cell-level TME descriptors add independent prognostic power to standard clinicopathological features.

Machine-vision tools such as those described here that produce interpretable image- and cell-level features can inform and improve current pathology practices, particularly in regard to stratification of risk, and have the potential to provide facile and scalable biomarkers for clinical studies in the future.

## ACKNOWLEDGEMENT

The results shown here are in whole or part based upon data generated by the TCGA Research Network: https://www.cancer.gov/tcga.

The authors would like to thank Patricia Spilman for her valuable feeback on the manuscript.

