## Supplementary material for "Deep learning-based risk stratification for HER2-negative breast cancer patients": Table S1, Table S2 and Table S3.

### *Supplementary information*

**Table S1** Features in baseline model introduced in Fig. 1a

| | **WSI** | TCGA-EW-A1P8-01Z-00-DX1 | TCGA-GM-A2D9-01Z-00-DX1 | TCGA-BH-A0B9-01Z-00-DX1 | TCGA-E2-A15I-01Z-00-DX1 | TCGA-AR-A24K-01Z-00-DX1 |
|---|---|---|---|---|---|---|
| **Clinicopathological features used in baseline model (Fig. 1a)** | *Prior treatment* | no | no | no | no | no |
| | *Ethnicity* | not hispanic or latino | not hispanic or latino | not reported | not hispanic or latino | not reported |
| | *Tissue of origin* | Breast, NOS | Breast, NOS | Breast, NOS | Breast, NOS | Breast, NOS |
| | *Treatment type* | Pharmaceutical Therapy, NOS | Radiation Therapy, NOS | Radiation Therapy, NOS | Pharmaceutical Therapy, NOS | Radiation Therapy, NOS |
| | *AJCC pathologic M* | M0 | M0 | M0 | M0 | M0 |
| | *Race* | white | white | white | white | white |
| | *Primary diagnosis* | Infiltrating duct carcinoma, NOS | Infiltrating duct carcinoma, NOS | Infiltrating duct carcinoma, NOS | Infiltrating duct carcinoma, NOS | Infiltrating duct carcinoma, NOS |
| | *AJCC pathologic N* | N3b | N0 (i-) | N0 (i-) | N0 | N0 |
| | *HR-status* | negative | positive | negative | positive | positive |
| | *AJCC pathologic T* | T2 | T1c | T1c | T2 | T2 |
| | *Prior malignancy* | no | no | no | no | no |
| **Predicted risk** | | low risk | low risk | high risk | high risk | high risk |

**Table S2** The most predictive features in clinicopathology- and image-based model

| **WSI** | **Top contributing features in clinicopathology- and image-based model (Fig. 6a)** | | | | | | **Predicted risk** |
|---|---|---|---|---|---|---|---|
| | *Prior treatment* | *Ethnicity* | *Tissue of origin* | *AJCC pathologic M* | *% sTALs* | *% stroma in tumor* | |
| TCGA-EW-A1P8-01Z-00-DX1 | no | not hispanic or latino | Breast, NOS | M0 | 47.07 | 1.46 | low risk |
| TCGA-GM-A2D9-01Z-00-DX1 | no | not hispanic or latino | Breast, NOS | M0 | 79.20 | 4.62 | low risk |
| TCGA-BH-A0B9-01Z-00-DX1 | no | not reported | Breast, NOS | M0 | 26.67 | 1.83 | high risk |
| TCGA-E2-A15I-01Z-00-DX1 | no | not hispanic or latino | Breast, NOS | M0 | 70.00 | 1.17 | low risk |
| TCGA-AR-A24K-01Z-00-DX1 | no | not reported | Breast, NOS | M0 | 59.67 | 18.51 | high risk |

**Table S3** The most predictive features in clinicopathology-, image-, and cell-based model

| **WSI** | **Top contributing features in clinicopathology-, image-, and cell-based model (Fig. 7a)** | | | | | | | | | | **Predicted risk** |
|---|---|---|---|---|---|---|---|---|---|---|---|
| | *Prior treatment* | *AJCC pathologic M* | *% sTALs* | *iTILs nucleus max caliper* | *iTILs nucleus eccentricity* | *Stroma in tumor nucleus max* | *Stroma in tumor nucleus* | *Stroma near tumor nucleus* | *Lymphocytes nucleus eccentricity* | *Lymphocytes nucleus max caliper* | |
| TCGA-EW-A1P8-01Z-00-DX1 | no | M0 | 47.1 | 5.17 | 0.76 | 4.96 | 52.0 | 216 | 0.76 | 5.17 | high risk |
| TCGA-GM-A2D9-01Z-00-DX1 | no | M0 | 79.2 | 5.14 | 0.79 | 5.02 | 52.0 | 221 | 0.82 | 5.18 | high risk |
| TCGA-BH-A0B9-01Z-00-DX1 | no | M0 | 26.7 | 5.28 | 0.77 | 5.11 | 52.2 | 280 | 0.77 | 5.30 | low risk |
| TCGA-E2-A15I-01Z-00-DX1 | no | M0 | 70.0 | 5.83 | 0.77 | 6.00 | 48.9 | 85.3 | 0.79 | 5.35 | low risk |
| TCGA-AR-A24K-01Z-00-DX1 | no | M0 | 59.7 | 5.33 | 0.77 | 5.20 | 51.7 | 166 | 0.78 | 5.34 | low risk |
